# A new species of the genus *Euphlyctis* (Anura: Dicroglossidae) from West Bengal, India

**DOI:** 10.1101/2021.04.18.440320

**Authors:** Somnath Bhakat, Soumendranath Bhakat

## Abstract

A new species of green frog of the genus *Euphlyctis* Fitzinger, 1843 is described from West Bengal, a state of eastern India. A robust frog, SVL of male 86mm and that of female 132mm. The species is diagnosed by the presence of following characters: green dorsum and female with a greenish white mid-dorsal line, tibiotarsal articulation reaches eye, male with two vocal sac openings at the junction of jaw, female is larger than male though body parameters is proportionately longer in male, nostril snout length 3.45% of SVL, nostril much closer to snout tip than eye, units of hind limb i. e. thigh, shank and foot are almost equal in length, relative length of finger: II < IV < I < III.

The new species is compared with existing eight species of the genus *Euphlyctis*.

## Introduction

In India, 284 species of amphibian are recorded of which only 50 species are reported from West Bengal (Dutta & Mukhopadhyay, 2013).The genus *Euphlyctis* comprising only eight currently recognized species: the most common *E. cyanophlyctis* (Schneider, 1799) from Iran, Afghanistan, Pakistan, Nepal, India, Sri Lanka, Malaya, and Vietnam; *E. hexadactylus* (Lesson, 1934) from India, Sri Lanka and Bangladesh; *E. ghoshi* (Chanda, 1991) from India (Manipur only); *E. karaavali* (Priti, Naik, Seshadri, Singal, Vidisha, Ravikanth & Gururaja, 2016) from India (Karnataka); *E. aloysii* (Joshy, Alam, Kurabayashi, Sumida & Kuramoto, 2009) from India (Mangalore); *E. mudigere* (Joshy, Alam, Kurabayashi, Sumida & Kuramoto, 2009) from India (Western Ghat) and *E. ehrenbergii* (Peters, 1863) from Saudi Arabia and Yemen and *E. kalasgramensis* (Howlader, Nair, Gopalan & Menla, 2015) from Bangladesh (Frost, 1985; Chanda, 1991; Dubois, 1992; Joshy et al., 2009; Howlader et al., 2015 and Priti et al., 2016).Among these two species viz. *E. cyanophlyctis* and *E. hexadactylus* are common in West Bengal (Srinivasulu et al., 2006). Recently I have collected a few species of frog from my locality (in and around Suri) by myself and with the help of other people. The present paper describes one such collected frog species of the genus *Euphlyctis, E. bengalensis* sp. nov.

## Materials and methods

Two live specimens of *Euphlyctis* (one male and one female) were collected at night from Bhadirban (six km. west of Suri), Birbhum district, West Bengal, India and kept in a large plastic jar with a few pores at the top. Next day, the specimens were killed by chloroform. Morphological data were taken with the help of a digital slide caliper, a divider and a millimeter scale. Photographs were taken immediately after killing. The specimens were preserved in 4% formaldehyde solution for further investigation.

The abbreviations used for morphometric analysis and comparison with other species are as follows: SVL= Snout vent length, HL= Head length (from the rear of mandible to the tip of snout), HW= Head width (length at the angle of jaws), HD= Head depth (vertical length posterior to orbit), UAL= Upper arm length (from the base of upper arm to elbow), FAL= Fore arm length (from the fixed elbow to the base of outer palmer tubercle), HAL= Hand length (from the of outer palmer tubercle to the tip of the third finger), THL= Thigh length (from the vent to the knee), SHL= Shank length (from knee to heel), FOL= Foot length (from the base of inner metatarsal tubercle to the tip of the fourth toe), TFOL= Distance from the heel to the tip of the fourth toe, TFLL= Total fore limb length, THLL= Total hind limb length, EN= Distance from the front of the eye to the nostril, EL= Eye length (horizontal distance between the orbital borders), SL= Snout length (from the tip of the snout to the anterior border), NS= Distance from the nostril to the tip of the snout, IN= Internarial distance (shortest distance in between two nostrils), IUE= Inter upper lid width (shortest distance between the upper eye lids), UEW= Maximum upper eye lid width, IFE= Interval front of the eyes (shortest distance between the anterior orbital borders), IBE= Interval back of the eyes (shortest distance between the post orbital borders), MN= Distance from the back of mandible to nostril, MFE= Distance from the back of mandible to the front of eye, MBE= Distance from the back of mandible to the back of eye, IOL= Inter orbital length, OTL= Posterior of orbit to anterior of tympanum length, TD= Tympanum diameter, FLI-IV= Finger length of I to IV, NP= Nuptial pad, TLI-V= Toe length of I to V, IMTL= Length of inner metatarsal tubercle, MTFF= Distance from distal edge of metatarsal to maximum incurvature of web between fourth and fifth toe, MTTF= Distance from distal edge of metatarsal to maximum incurvature of web between third and fourth toe, TFTF= Distance from maximum incurvature of web between third and fourth toe to tip of fourth toe, FFTF= Distance from maximum incurvature between fourth and fifth toe to tip of fourth to tip of fifth toe.

*E. bengalensis* was compared with other eight species of *Euphlyctis* on the basis of different morphometric characters by paired t-test. Spearman’s correlation was performed to separate different morphometric characters of male and female frogs. Different body parameters (HL, HW, etc.) of nine species of *Euphlyctis* were compared by using ANOVA to find out if the mean values of different body parameters are significantly different or not.

Principal component analysis (PCA) and construction of heatmap was performed on the multivariate data of nine species of *Euphlyctis* (see Table S1 in Supplementary Informations). PCA and heatmap analyses were performed using ClustVis: https://biit.cs.ut.ee/clustvis/ (Metsalu & Vilo, 2015). PCA does a linear transformation on the multivariate data to extract variables (principal components) that are uncorrelated. The first few components (otherwise known as principal components and denoted as PC1, PC2) capture the data with most variability. Plotting PC1 *vs* PC2 approximate the distance between points and if the points on the 2D PC plots are not overlapping then these points form separate clusters. Before PCA analyses, the original values are transformed using *ln(x+1)* function. Unit variance scaling was applied on the row values of the dataset and *singular value decomposition* (SVD) with imputation was used to calculate PCs.

Heatmap uses color coding to visualize multivariate data matrix. Looking at the phenogram integrated with heatmap one can get a *helicopter* view of how corresponding variables of different species are separated better than others.

## Result

### Taxonomy

Amphibia Linnaeus, 1758

Anura Fisher von Waldhein, 1813

Dicroglossidae Anderson, 1871

Dicroglossinae Anderson, 1871

*Euphlyctis* Fitzinger, 1843

*Euphlyctis bengalensis* sp. nov. (Fig. 1).

**FIGURE 1.**
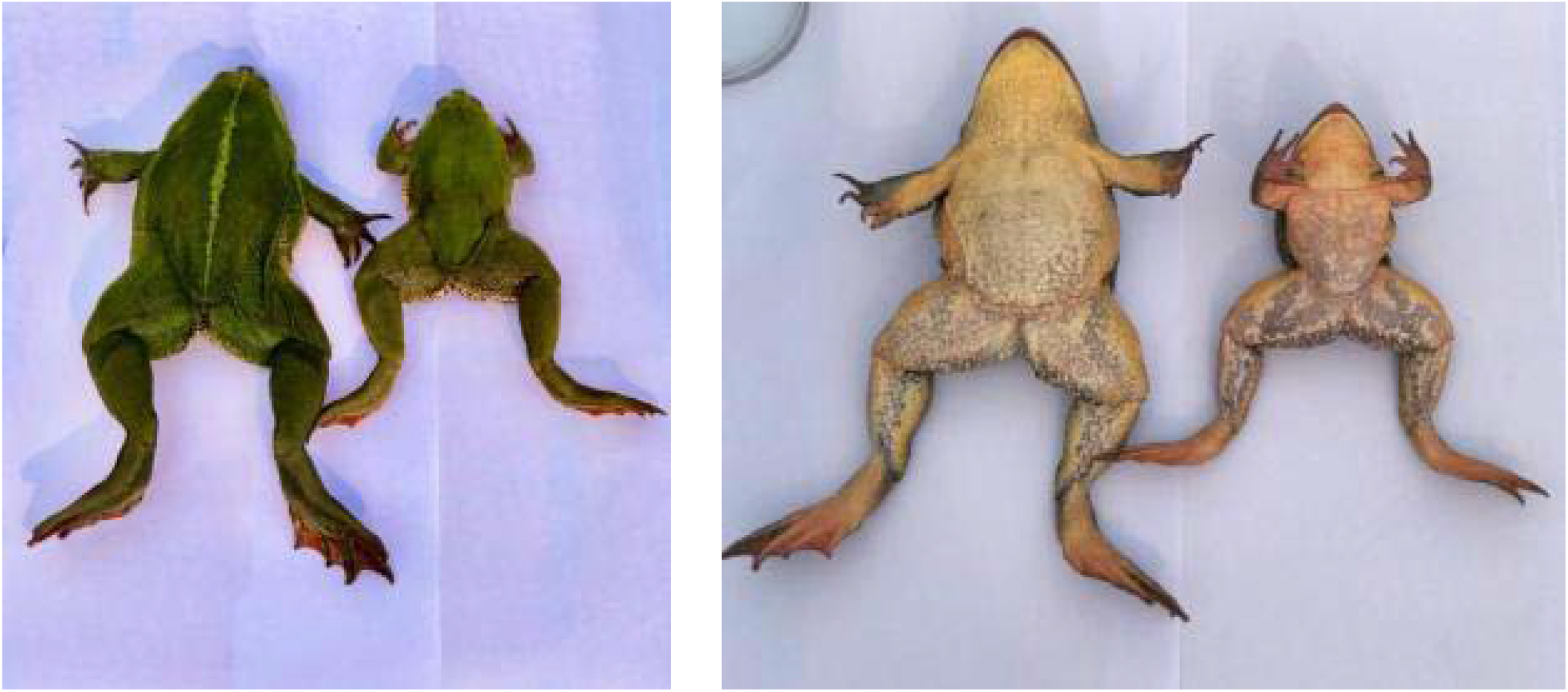
*Euphlyctis bengalensis*. Adult female and male: Dorsal view (left), ventral view (right).

#### Holotype

Zoological Museum, Department of Zoology, Rampurhat College, Rampurhat-731224, Dist. Birbhum, W. B., India, mature male, SVL: 86 mm. (**Fig. 1**), collected in Bhandirban, six km. west of Suri (87°32’00’’E, 23°55’00’’N) Birbhum District, on 8^th^ December 2020, by Anshu Rajak,

#### Paratype

Mature female, SVL: 132 mm (**Fig. 1**), other information same as holotype.

#### Etymology

Species name is an adjective to the ‘bengal’, the state (West Bengal) from where holotype was collected.

#### Diagnosis

Large and robust *Euphlyctis* species. The species is characterized by a combination of the following characters: SVL of holotype male 86 mm and that of paratype female 132 mm, dorsal dark green with a greenish white middorsal line in female only, nostril snout length 3.45% of SVL (n= 2), nostril much closer to snout tip than eye (more than double in length), nostril snout length 42.77% of distance from eye to nostril, fore arm length 18.89% of SVL, foot length 48.81% of SVL. Relative length of fingers: II< IV< I< III. Thigh, shank and foot are almost equal in length in both sexes. Relative length of toes: I< II< III< V< IV.

#### Description

(Table 1).Adult male (Holotype): Large sized frog (SVL 86 mm). Head large, conical, broader than long (HW: HL= 1.25), a little depressed, canthus rostralis indistinct, loreal region oblique and concave, snout obtusely pointed both dorsally and ventrally, projecting beyond mouth. In lateral view, snout bluntly pointed. Snout length equal to eye length (EL= SL= 12 mm). Nostril much closer to snout than eyes (NS= 3.0, EN= 7.4), round and covered with a small flap (**Fig. 3**). Internarial distance slightly greater than interorbital length (IN= 4.3 mm, IOL= 4.0 mm) which is half of eye length (EL= 8 mm). Eye is moderately large (EL 28.57% of HL and 9.30% of SVL), protruding, pupil horizontal, iris blackish. Upper eye lid width is 1.5 times greater than interorbital length (UEW 21.43% of HL, IOL 14.29% of HL). Tympanum is round, large and close to eye (TD 30% of HL, OTL 10.99% of HL) and slightly larger than eye (EL 95% of TD). Supratympanic fold distinct. A distinct fold of skin runs from posterior of orbit to the base of shoulder covering the tympanum (**Fig. 3**). A pair of vocal sac openings on both sides of lower jaw near jaw angle in the form of 8 mm long slit (**Fig. 4**). The slit is as long as the length of eye. Vomerine teeth are oblique in position, in the anterior margin of choanae. Spatuate bifid tongue is with lingual papilla (**Fig. 4**). Lower jaw with three tooth like bony projections that fit in fossa on the upper jaw (**Fig. 2**).

**TABLE. 1.** Morphometric data of *Euphlyctis bengalensis* sp. nov. (measurements in mm.).

| Sex |  | Male |  | Female |  |  |  |  | Male |  | Female |  |  |
| --- | --- | --- | --- | --- | --- | --- | --- | --- | --- | --- | --- | --- | --- |
| Sl. No. | Characters | Values | % SVL | Values | % SVL | Mean | Sl. No. | Characters | Values | % HL | Values | % HL | Mean |
| 1. | SVL | 86 |  | 132 |  |  | 23. | MN | 23.6 | 84.29 | 34 | 89.47 | 86.88 |
| 2. | HL | 28 | 32.56 | 38 | 28.79 | 30.68 | 24. | MFE | 17.5 | 62.50 | 25 | 65.79 | 64.15 |
| 3. | HW | 35 | 40.70 | 47.7 | 36.14 | 38.42 | 25. | MBE | 12.5 | 44.64 | 17.5 | 46.05 | 45.35 |
| 4. | HD | 14.5 | 16.86 | 21 | 15.91 | 16.39 | 26. | IOL | 4 | 14.29 | 5.2 | 13.68 | 13.99 |
| 5. | UAL | 19 | 22.09 | 21 | 15.91 | 19.00 | 27. | OTL | 3.5 | 12.50 | 3.6 | 9.47 | 10.99 |
| 6. | FAL | 17.5 | 20.35 | 23 | 17.42 | 18.89 | 28. | TD | 8.4 | 30.00 | 10 | 26.32 | 28.16 |
| 7. | HAL | 23 | 26.74 | 28.5 | 21.59 | 24.17 |  |  |  | % TFL |  | % TFL |  |
| 8. | THL | 45 | 52.33 | 60 | 45.45 | 48.89 | 29. | FLI | 17.5 | 29.41 | 24 | 33.10 | 31.26 |
| 9. | SHL | 45 | 52.33 | 58.5 | 44.32 | 48.33 | 30. | FLII | 15.5 | 26.05 | 20.5 | 28.28 | 27.17 |
| 10. | FOL | 45.5 | 52.91 | 59 | 44.70 | 48.81 | 31. | FLIII | 20.5 | 34.45 | 28.2 | 38.90 | 36.68 |
| 11. | TFOL | 69.5 | 80.81 | 90 | 68.18 | 74.50 | 32. | FLIV | 16.5 | 27.73 | 22 | 30.34 | 29.04 |
| 12. | TFL | 59.5 | 69.19 | 72.5 | 54.92 | 62.06 | 33. | NP | 4.2 | 7.06 | - | - | - |
| 13. | THLL | 149.5 | 173.84 | 208.5 | 157.95 | 165.90 |  |  |  | % THLL |  | % THLL |  |
|  |  |  | % HL |  | % HL |  | 34. | TLI | 19.5 | 13.04 | 19.5 | 9.35 | 11.20 |
| 14. | EN | 7.4 | 26.43 | 10 | 26.32 | 26.38 | 35. | TLII | 25.5 | 17.06 | 30 | 14.39 | 15.73 |
| 15. | EL | 8 | 28.57 | 12 | 31.58 | 30.08 | 36. | TLIII | 31.5 | 21.07 | 41.5 | 19.90 | 20.49 |
| 16. | SL | 8 | 28.57 | 12 | 31.58 | 30.08 | 37. | TLIV | 40.2 | 26.89 | 54.2 | 26.00 | 26.45 |
| 17. | NS | 3 | 10.71 | 4.5 | 11.84 | 11.28 | 38. | TLV | 34.2 | 22.88 | 45 | 21.58 | 22.23 |
| 18. | IN | 4.3 | 15.36 | 5 | 13.16 | 14.26 | 39. | IMTL | 3.8 | 2.54 | 5.5 | 2.64 | 2.59 |
| 19. | IUE | 16 | 57.14 | 18.5 | 48.68 | 52.91 | 40. | MTFF | 28.5 | 19.06 | 35 | 16.79 | 17.93 |
| 20. | UEW | 6 | 21.43 | 9 | 23.68 | 22.56 | 41. | MTTF | 32 | 21.40 | 39 | 18.71 | 20.06 |
| 21. | IFE | 13.2 | 47.14 | 16.5 | 43.42 | 45.28 | 42. | TFTF | 11.3 | 7.56 | 14 | 6.71 | 7.14 |
| 22. | IBE | 18.5 | 66.07 | 25.5 | 67.11 | 66.59 | 43. | FFTF | 15 | 10.03 | 20 | 9.59 | 9.81 |

**FIGURE 2.**
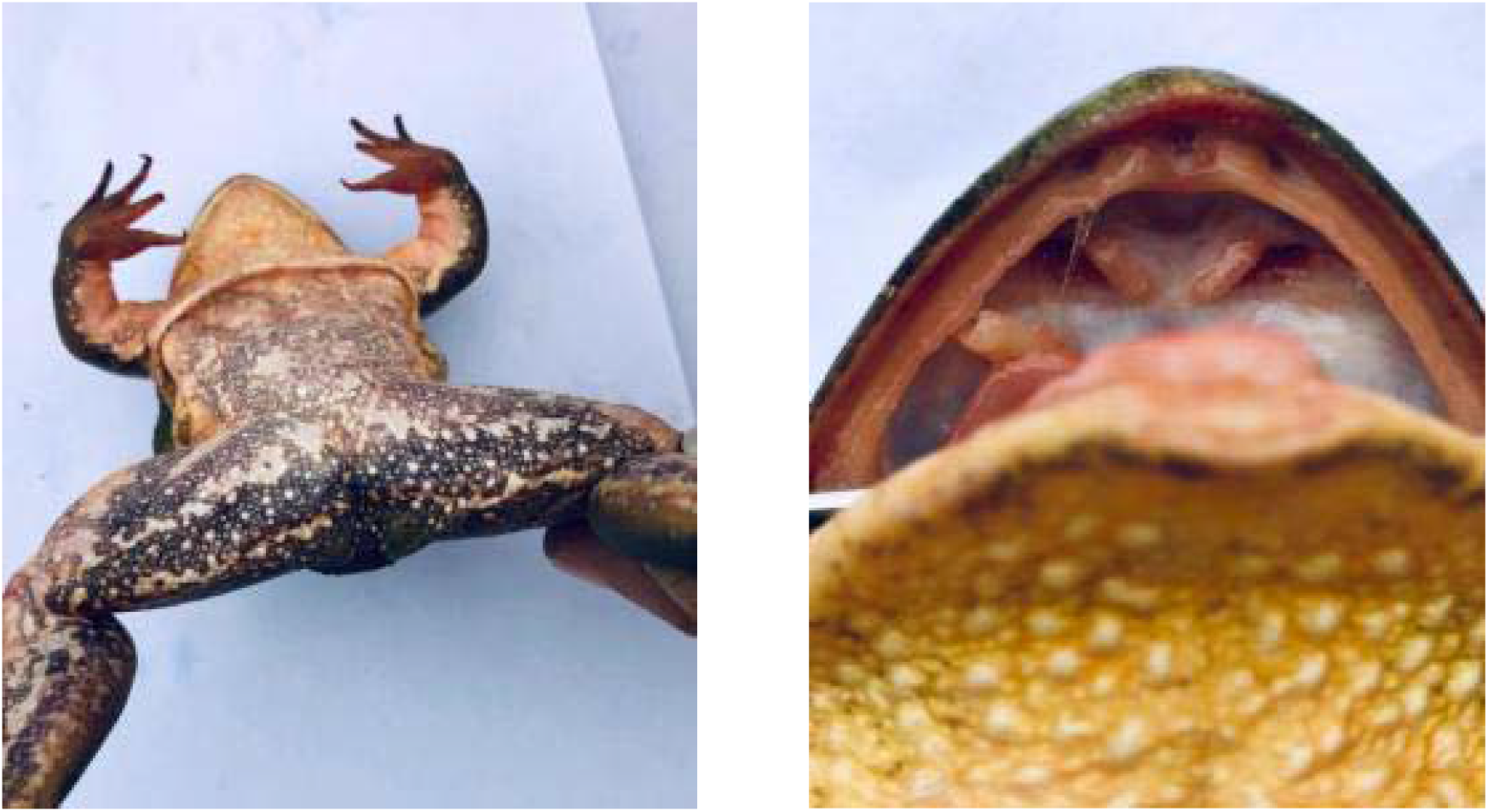
*Euphlyctis bengalensis*. Posterior of thigh (left), interior of mouth showing fossa on upper jaw (right).

**FIGURE. 3.**
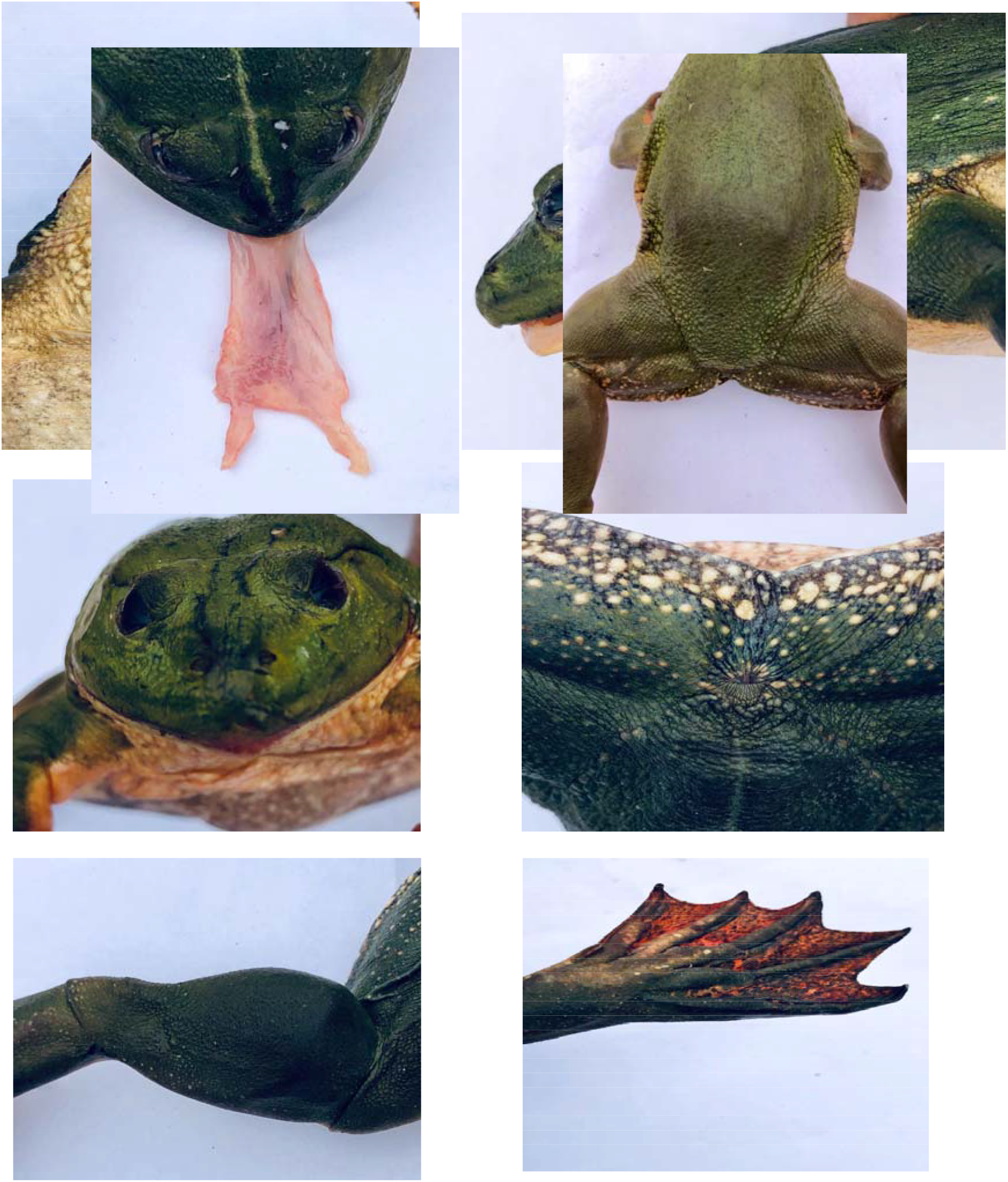
*Euphlyctis bengalensis*. Throat showing warts (upper left), skin fold from orbit to shoulder and tympanum (upper right), anterior view of head (middle left), vent (middle right), knee and ankle cap (lower left), web and toes (lower right).

**FIGURE. 4.**
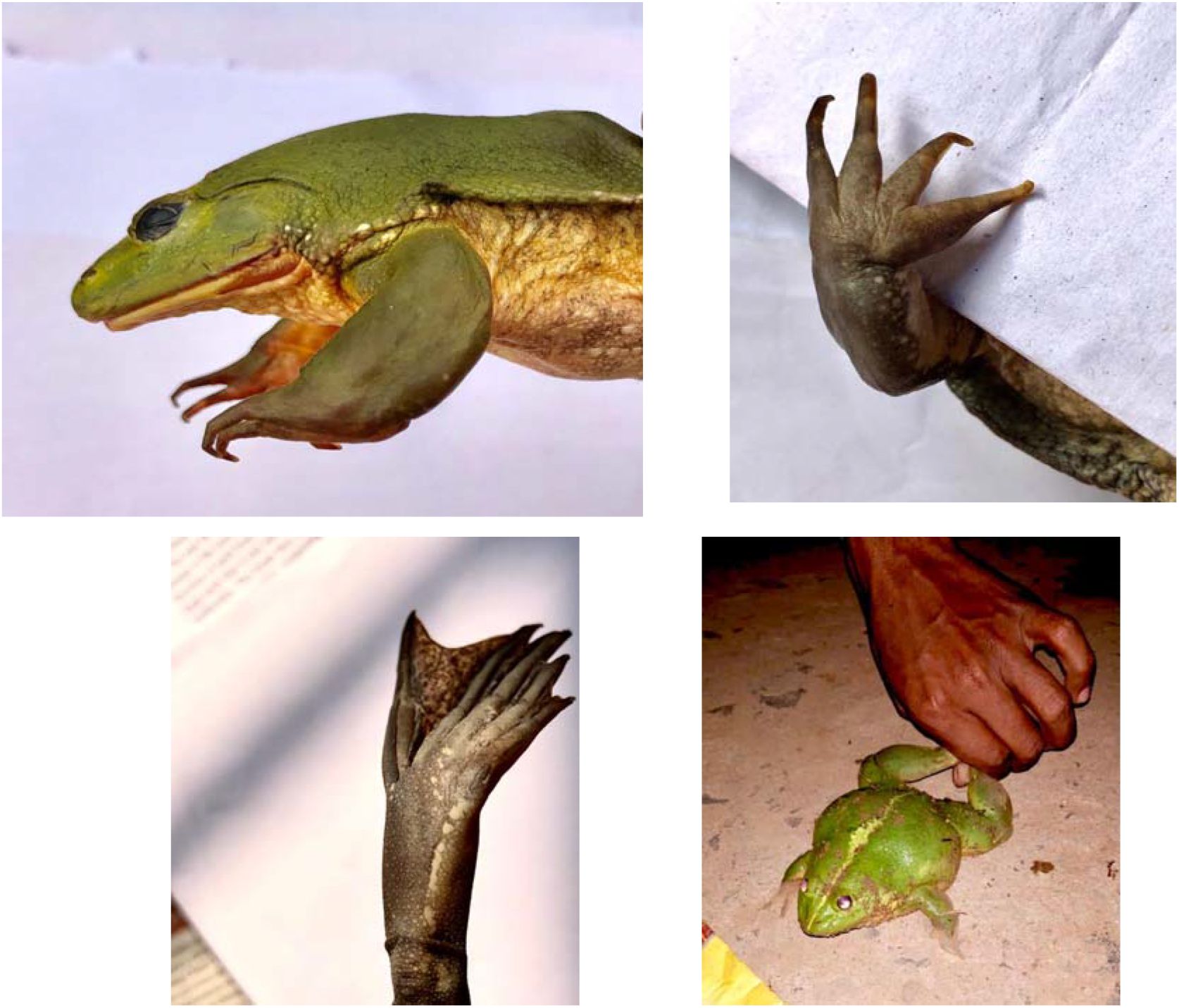
*Euphlyctis bengalensis*. Bifid tongue (upper left), ‘U’ shaped structure between thighs (upper right), vocal sac opening in male (middle left), finger (middle right), white spotted line in foot (lower left), a live female (lower right).

##### Forelimb

Strong and stout, 69.19% of SVL. Fore arm length 1.3 times the hand length (FAL= 17.5 mm, HAL= 23 mm), fingers free, gradually tapering to pointed tips though base if first finger is very wide (**Fig. 4**). First finger is longer than the second. Red blister like oval nuptial pad (4.2 mm long and 3.1 mm width) is at the base of first finger (according to Flores, 1985, nuptial pad is Type I). Second and third fingers slightly fringed laterally. Subarticular tubercles are small and indistinct. Relative lengths of fingers: II< IV< I< III (FLI= 17.5 mm, FLII= 15.5 mm, FLIII= 20.5 mm, FLIV= 16.5 mm).

##### Hindlimb

Long and stout, 1.74 times of SVL. Heels overlap when folded at right angles to body, tibiotarsal articulation reaching eyes. Thigh, shank and foot length almost equal in length and more than half of SVL (52% of SVL) with cap like skin fold in knee and ankle (**Fig. 3**). Heel to tip of fourth toe 1.7 times longer than fourth toe length (TFOL= 69.5 mm, TLIV= 40.2 mm). Inner metatarsal tubercle is small and digitiform, about one-fifth of first toe length (IMTL= 3.8 mm, TLI= 19.5 mm), outer metatarsal tubercle absent. Toe tip small and slightly pointed. Subarticular tubercles are small and indistinct. Basal half of each toe is broad with skin folds while distal half is narrow (Fig. 3). First and fifth toe with well develop fringe (wider in the first toe). Relative toes lengths: I< II< III< V< IV (TLI= 19.5 mm, TLII= 25.5 mm, TLIII= 31.5 mm, TLIV= 40.2 mm, TLV= 34.2 mm). Fully developed web reaching toe tip and incised sharply, MTTF greater than MTFF (1.12 times) and FFTF is also greater than TFTF (1.33 times).

##### Adult female (Paratype)

Most of the body parameters in female are proportionately lower compared to those of male. Both the fore and hind limbs in female is shorter than those of male (26% and 10% respectively). Moreover, finger and toe length in percentage of SVL is longer in male compared to those of female. Webs of hind limbs are also well developed in male. Some characters exhibited significant differences between sexes though proportionate values are within a reasonable range. 18 characters (HL, HW, UAL, FAL, HAL, THL, SHL, FOL, TFOL, EL, SL, IUE, IFE, MN, OTL, TD, FLI and TLI) of male and female frog are significantly different (Spearman’s correlation, r_s_= 0.35).

##### Skin

Dorsum with small warts mostly on the posterior one-third and lateral side of the body. Fore limb smooth while hind limb with small warts except from heel to toes. A ‘U’-shaped line of warts from upper portion of thigh to vent is present in both sexes (**Fig. 4**). Ventral is with porous warts, more distinct and large on the thorax and posterior of thigh.

##### Colour (in live)

Dorsal leafy green, female with a greenish white mid-dorsal line from vent to snout tip (**Fig. 1** and **4**), a few white spot on the lateral side of the body and upper portion of hand. Tympanum is also dark green and difficult to separate from body colour (**Fig. 3**). A few white spots are present at the base of hand and leg. Sides of body from hand to leg and lower part of the dorsum (in between two thighs) are with distinct green warts. Ventral portion of 1st and 2^nd^ finger are greenish while 3^rd^ and 4^th^ finger yellowish red. 1^st^ 2^nd^ and 3^rd^ toes reddish white while 4^th^ and 5^th^ toes are green. A white spotted line is present from ankle to base of first toe (Fig. 4). Ventral is yellowish in female and yellowish brown in male. In female, brownish patches are present particularly in thigh, shank and lower half of abdomen. Yellowish white large round warts are distinct on throat and breast (**Fig. 3**). Brownish ‘V’-shaped patch with white spot in thigh and shank is present in male (pale in female) (**Fig. 1**). Abdomen also have pale brownish patch. Posterior of thigh is brownish in colour with white irregular roundish spots (Fig. 2). Greenish narrow fold of skin is present around the vent (**Fig. 3**). Tongue reddish. Web bears dark marblings (**Fig. 3**).

#### Natural history

*E. bengalensis* sp. nov. a semi aquatic frog usually live half submerged in water throughout the day but at night often come to land for roaming and collection of food. They are very common in ponds and wetlands with full of water hyacinth and other species of floating plants.

#### Distribution

Presently known only from the type locality, Bhandirban, a small village, six km. west of Suri, Dist. Birbhum, West Bengal, India.

#### Comparison

Comparisons of the new species with other eight species of *Euphlyctis* was done using data from literature (Boulenger, 1920; Chanda, 1990; Joshy et. al., 2009; Howlader et al., 2015; and Priti et al., 2016). The present species, *E. bengalensis* sp. nov. distinguished from all other existing species of *Euphlyctis* on several aspects like distribution, colour, size and most importantly on morphology. Except four green *Euphlyctis*, viz. *E. hexadactylus, E. karaavali, E. ehrenbergii* and the present species *E. bengalensis* sp. nov.; dorsum of all the five species is light brown to grayish brown with irregular spots. Moreover, all three Indian species are larger in size than rest of the species and males are distinctly smaller than females. Maximum SVL of male and female are 93 mm. and 130 mm. respectively in *E. hexadactylus* (Boulenger, 1920), 86 mm. and 132 mm. respectively in *E. bengalensis* (present study). In *E. karaavali*, males and females are comparatively smaller (male: 61.89 mm. and female: 106.3 mm.) (Priti et al., 2016).

The present species is clearly distinguished from its closest congener *E. hexadactylus* in having a smaller ratio of HL/HW (0.799 vs. 0.988), SN/NE (0.428 vs. 0.878), NN/EE (1.019 vs. 1.465) and larger ratio of TD/ED (0.94 vs. 0.87), FI/FII (1.15 vs. 1.11). NS in the present species is less than half of that of *E. hexadactylus* (3.45% SVL vs. 8.10% SVL). Snout in *E. hexadactylus* is longer compared to *E. bengalensis* (11.35% SVL vs. 20% SVL). In *E. hexadactylus* IOD is 23% less and TD is more than 10% greater compared to *E. bengalensis*. Moreover, all the fingers and toes are proportionately longer in *E. bengalensis*. In *E. hexadactylus* a prominent black central stripe from base of fore limb is present vs. absent in the present species. *E. hexadactylus* was reported to have a white or pale yellow venter (Dutta & Manamendra-Arachchi, 1996; Chanda, 2002; Daniel, 2002; Daniels, 2005) and the specimens of Mangalore have a finely mottled pattern on the venter (Joshy et al., 2009). But in the present species, venter is dark green with a few white spots around it.

*E. bengalensis* differs from *E. karaavali* in having higher ratio of ELW/EE (1.616 vs. 1.382) and smaller ratio of SN/NE (0.428 vs. 0.706) and HL/HW (7.99 vs. 0.935). Further snout of *E. karaavali* is 1.78 times longer than that of *E. bengalensis*. HL, EL, NS and IN percentage of SVL is higher in *E. karaavali* compared to *E. bengalensis* (35.26 vs. 30.68, 10.30 vs. 9.20, 6.67 vs. 3.45 and 5.56 vs. 4.40 respectively) while reverse is true for SHL and THL (44.81 vs. 48.83 and 43.16 vs. 48.89 respectively).

*E. bengalensis* is distinguished from *E. kalasgramensis* of Bangladesh by higher ratio of TD/ED (0.940 vs. 0.626) and smaller ratio of NN/EE (1.019 vs. 1.256). Snout and eye length of *E. kalasgramensis* are 20% and 69% greater than those of *E. bengalensis* while units of hind limb (THL, SHL, FOL) is more than 20% greater compared to the present species. Head is longer in length in the species from Bangladesh (33.15% SVL vs. 30.68% SVL).

NS and IN of *E. aloysii* is 1.85 and 1.5 times greater than those of *E. bengalensis*. Snout of *E. aloysii* is 17% longer than that of *E. bengalensis* though HW and THL (in % of SVL) are shorter (33.00 vs. 38.42 and 43.90 vs. 48.89). Further, *E. aloysii* differs from *E. bengalensis* in having higher ratio of SN/NE, NN/EE (0.925 vs. 0.428, 1.569 vs. 1.019) and smaller ratio of FI/HAL (0.607 vs. 0.801). Moreover, fingers and toes of *E. bengalensis* are longer than those of *E. aloysii*. In *E. bengalensis* THL, SHL and FOL are equal in length while THL is comparatively smaller in *E. aloysii* (SHL = FOL > THL).

Nostril to snout distance is more than double in *E. mudigere* than those of *E. bengalensis* (7.70% SVL vs. 3.45% SVL). Moreover, EL and IN percentage of SVL is greater in *E. mudigere* (12.00 vs. 9.20 and 6.90 vs. 4.40). *E. bengalensis* differs from *E. mudigere* in having smaller ratio of SN/NE and NN/NE (0.428 vs. 1.124 and 1.019 vs. 1.505) and higher ratio of HW/SVL and HAL/SVL (38.42 vs. 35.50 and 24.17 vs. 21.90). Further fingers and toes of *E. bengalensis* are longer than those of *E. mudigere*.

In *E. bengalensis* all three parts of hind limb, THL, SHL and FOL are equal in length while in *E. cyanophlyctis* though THL and FOL are equal but SHL is greater in length (48% of SVL in *E. bengalensis*, SHL in *E. cyanophlyctis* 50% of SVL). *E. bengalensis* differs from *E. cyanophlyctis* in having smaller ratio of HL/HW (0.799 vs. 0.927), SN/NE (0.428 vs. 0.952) and NN/NE (1.019 vs. 1.451). Snout and eye in *E. cyanophlyctis* is 28% and 41% longer compared to those of *E. bengalensis*. Moreover, the values of NS, IN and TD in percentage of SVL is greater in *E. cyanophlyctis* (7.50 vs. 3.45, 6.10 vs. 4.40 and 11.30 vs. 8.68 respectively). Except first finger all other fingers and toes are slightly longer in the present species.

*E. ghoshi* not only restricted to Manipur but also very distinct from other species of *Euphlyctis* by combination of characters including rounded head, snout length 15% of SVL, tympanum diameter 6.67% of SVL and with minimum TD/ED ratio (only 0.50). Except the recently discovered *E. karaavali*, snout length in percentage of SVL is minimum in all other species (< 15%).

*E. ehrenbergii* is a non-Indian species and resembles with the Indian species, *E. hexadactylus*. It differs from the present species, *E. bengalensis* in having smaller ratio of TD/ED (0.596 vs. 0.940), NN/EE (0.84 vs. 1.019), ELW/EE (1.364 vs. 1.616), TD/SVL (6.80 vs. 8.68) and greater ratio of HAL/SVL (30.00 vs. 24.17), NS/SVL (7.00 vs. 3.45) and EL/SVL (11.50 vs. 9.20).

Pair-wise comparisons of *E. bengalensis* with other eight species of *Euphlyctis* are presented in **Table 2**. The result shows that except two foreign species, *E. kalasgramensis* (from Bangladesh) and *E. ehrenbergi* (from Saudi Arabia) all six Indian species are significantly different from the present species.

**TABLE 2.** Statistics obtained from pair wise comparison using measurements of nine species of *Euphlyctis*. Abbreviations: beng.= *E. bengalensis*, kara.= *E. karaavali*, ghos.= *E. ghoshi*, kala.= *E. kalasgramensis*, aloy.= *E. aloysii*, mudi.= *E. mudigere*, cyan.= *E. cyanophlyctis*, hexa.= *E. hexadactylus*, ehre.= *E. ehrenbergi*.

| Sl. No. | Species compared | No. of variables | t value | P |
| --- | --- | --- | --- | --- |
| 1. | beng. vs. kara. | 25 | 2.73 | <0.01 and 0.05 |
| 2. | beng. vs. ghos. | 12 | 2.10 | <0.05 |
| 3. | beng. vs. kala. | 17 | 1.62 | Not significant |
| 4. | beng. vs. aloy. | 22 | 3.06 | <0.01 and 0.05 |
| 5. | beng. vs. mudi. | 22 | 3.37 | <0.01 and 0.05 |
| 6. | beng. vs. cyan. | 23 | 4.07 | <0.01 and 0.05 |
| 7. | beng. vs. hexa. | 23 | 3.98 | <0.01 and 0.05 |
| 8. | beng. vs. ehre. | 13 | 0.79 | Not significant |

Result of ANOVA shows that different body parameters are not significantly different in nine species of *Euphlyctis* including the present species (Table 3).

**TABLE 3.** Calculations of ANOVA (body parameters of nine species of *Euphlyctis*).

| Due to | Degrees of freedom (df) | S. S. | M. S. | F- value |
| --- | --- | --- | --- | --- |
| Species | 8 | 542.47 | 67.81 | 0.296 |
| Error | 160 | 36663.11 | 229.14 |  |

PCA of multivariate (12 variables as shown in Supplementary Informations) data of nine species of Euphlyctis is presented in **Fig. 5**. PC1 and PC2 capture 63% variance in our data. Projection of PCs along first two components gave us an indication that the morphological variables of *E. hexadactylus* and *E. cyanophlyctis* are closer to each other. PCs clearly shows that based on morphological features, *E. bengalensis* is a distinct species and separated from all the other species of Euphlyctis.

**FIGURE 5.**
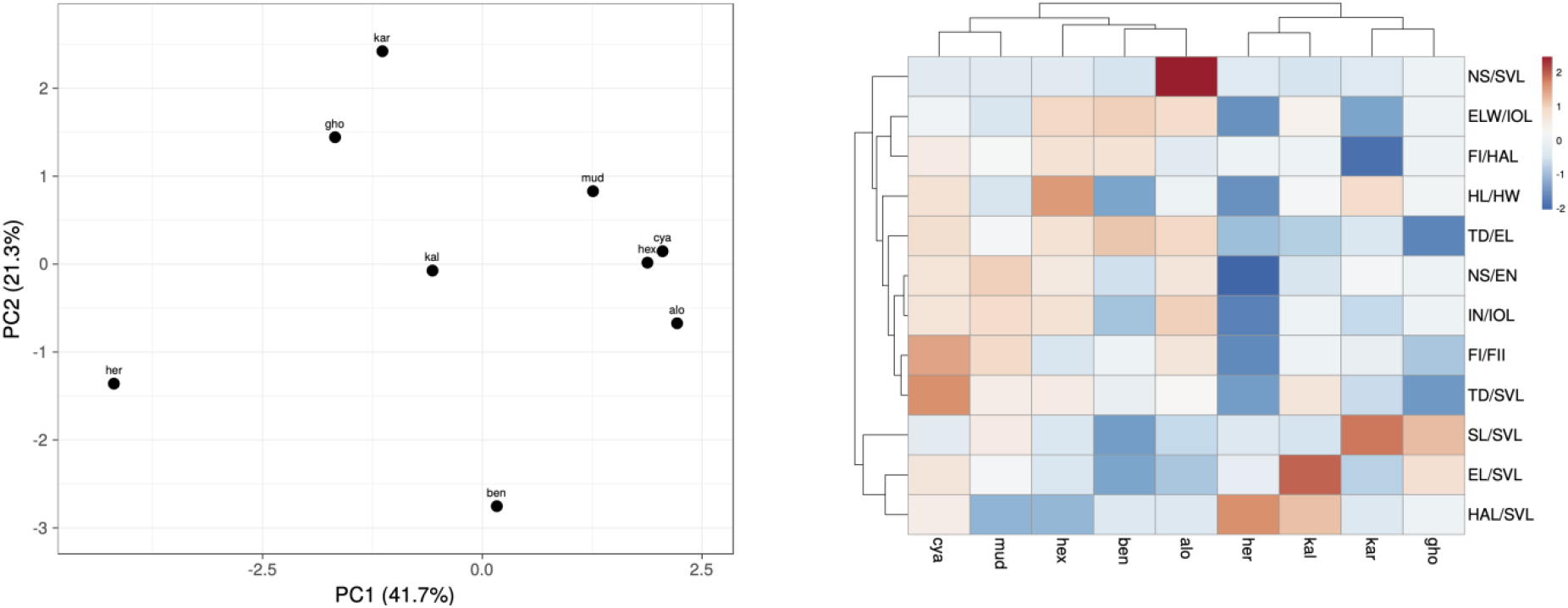
PCA was performed using ClustVis online interface. First two principal components capture maximum variance in the data and shows separation of different species. Heatmap and phenogram shows how different morphological variables of different species are grouped with each other.

Heat map including phenogram is presented in **Fig. 5**. Phenogram shows that for some morphological features E. bengalensis is morphologically more closure to *E. aloysii* however further genetic studies are necessary to compare these species beyond morphological characteristics.

## Discussion

The type locality of all the Indian species of *Euphlyctis* are south and south-western part of India (Boulenger, 1920; Daniels, 1992; Biju, 2001; Joshy, 2009; Priti et al. 2016) except *E. ghosi*, the type locality of which is Manipur only (Chanda, 1990). Among these, only two species, *E. cyanophlyctis* and *E. hexadactylus* are also reported from eastern part of India including West Bengal (Srinivasulu et al., 2006). So far eastern part of India especially West Bengal have not been explored fully for amphibian species. In this context, this is the first report of new discovery of an amphibian species of the genus *Euphlyctis* from West Bengal.

The discovery of the present species from pond of agricultural land indicates that there is high diversity of amphibian species outside of forest. In this region, random use of chemical fertilizer and pesticides in agriculture is the major cause of dwindling of different animal species including amphibians. To explore the animal species in any locality, one of the best ways is to engage common man of that locality. On this principle, I have already started to explore different animal species by citizen engagement in my locality (Birbhum district, West Bengal) and reported four new species of freshwater fish and a new species of land planarian, *Bipalium bengalensis*. Moreover, I have collected a few species of frogs (of the genus *Euphlyctis, Polypedates, Microhyla* and *Fejervarya*), millipedes (Polydesmida and Harpagophorida), dipteran fly and aquatic insects. The present report is an outcome of this drive.

## Acknowledgement

I am deeply indebted to Mr. Anshu Rajak of Bhandirban for collection of specimens and Mr. Samriddha Sil for technical help. I am also grateful to all my colleagues for their cooperation and finally to my wife for inspiring me constantly during the study period.

## Supplementary Informations

**TABLE S1.** Different morphological features corresponding to nine different species of *Euphlyctis*. N/A denotes data unavailability.

|  | **ben** | **her** | **cya** | **kal** | **mud** | **hex** | **alo** | **kar** | **gho** |
| --- | --- | --- | --- | --- | --- | --- | --- | --- | --- |
| **HL/HW** | 0.799 | 0.785 | 0.927 | 0.889 | 0.849 | 0.98 | 0.877 | 0.935 | 0.886 |
| **NS/EN** | 0.428 | N/A | 0.952 | 0.467 | 1.124 | 0.878 | 0.925 | 0.706 | N/A |
| **TD/EL** | 0.94 | 0.596 | 0.89 | 0.626 | 0.773 | 0.873 | 0.913 | 0.689 | 0.5 |
| **IN/IOL** | 1.019 | 0.84 | 1.451 | 1.256 | 1.505 | 1.465 | 1.569 | 1.091 | N/A |
| **ELW/IOL** | 1.616 | 1.364 | 1.515 | 1.549 | 1.466 | 1.607 | 1.601 | 1.382 | N/A |
| **FI/HAL** | 0.801 | N/A | 0.76 | N/A | 0.693 | 0.803 | 0.607 | 0.334 | N/A |
| **FI/FII** | 1.15 | 1 | 1.295 | N/A | 1.24 | 1.109 | 1.213 | 1.144 | 1.067 |
| **EL/SVL** | 0.092 | 0.115 | 0.13 | 0.156 | 0.12 | 0.11 | 0.1 | 0.103 | 0.133 |
| **TD/SVL** | 0.087 | 0.068 | 0.113 | 0.098 | 0.095 | 0.096 | 0.092 | 0.079 | 0.067 |
| **SL/SVL** | 0.092 | 0.115 | 0.118 | 0.111 | 0.134 | 0.114 | 0.108 | 0.164 | 0.15 |
| **HAL/SVL** | 0.242 | 0.3 | 0.265 | 0.286 | 0.219 | 0.221 | 0.241 | 0.241 | N/A |
| **NS/SVL** | 0.035 | 0.07 | 0.075 | 0.032 | 0.077 | 0.081 | 0.64 | 0.067 | N/A |

